# Genetic variation within genes associated with mitochondrial function is significantly associated with later age at onset of Parkinson disease and contributes to disease risk

**DOI:** 10.1101/475111

**Authors:** Kimberley J. Billingsley, Ines A. Barbosa, Sara Bandrés-Ciga, John P. Quinn, Vivien J. Bubb, Charu Deshpande, Juan A. Botia, Regina H. Reynolds, David Zhang, Michael A. Simpson, Cornelis Blauwendraat, Ziv Gan-Or, J Raphael Gibbs, Mike A. Nalls, Andrew Singleton, International Parkinson’s Disease Genomics Consortium (IPDGC), Mina Ryten, Sulev Koks

## Abstract

Mitochondrial dysfunction has been implicated in the aetiology of monogenic Parkinson’s disease (PD). Yet the role that mitochondrial processes play in the most common form of the disease; sporadic PD, is yet to be fully established. Here we comprehensively assessed the role of mitochondrial function associated genes in sporadic PD by leveraging improvements in the scale and analysis of PD GWAS data with recent advances in our understanding of the genetics of mitochondrial disease. First, we identified that a proportion of the “missing heritability” of the PD can be explained by common variation within genes implicated in mitochondrial disease (primary gene list) and mitochondrial function (secondary gene list). Next we calculated a mitochondrial-specific polygenic risk score (PRS) and showed that cumulative small effect variants within both our primary and secondary gene lists are significantly associated with increased PD risk. Most significantly we further report that the PRS of the secondary mitochondrial gene list was significantly associated with later age at onset. Finally, to identify possible functional genomic associations we implemented Mendelian randomisation, which showed that 14 of these mitochondrial function associated genes showed functional consequence associated with PD risk. Further analysis suggested that the 14 identified genes are not only involved in mitophagy but implicate new mitochondrial processes. Our data suggests that therapeutics targeting mitochondrial bioenergetics and proteostasis pathways distinct from mitophagy could be beneficial to treating the early stage of PD.

## INTRODUCTION

Parkinson’s disease (PD) is a progressive neurodegenerative movement disorder characterized pathologically by the death of dopaminergic neurons in the substantia nigra (SN) and aggregation of α-synuclein protein (encoded by *SNCA*), within intraneuronal inclusions called Lewy bodies. The majority of PD cases are apparently sporadic in nature (∼95%). Aging is a major risk factor for the disease and due to population ageing the prevalence of PD is predicted to increase rapidly, making the identification of therapeutic targets a high priority^1–3^.

Although there have been great advances in understanding both the genetic architecture and cellular processes involved in PD, the exact molecular mechanisms that underlie PD remain unknown ^1^. However, it is suggested that PD has a complex etiology, involving several molecular pathways, and understanding these specific pathways will be key to establishing mechanistic targets for therapeutic intervention. While several key pathways are currently being investigated, including autophagy, endocytosis, immune response and lysosomal function, ^4–7^ mitochondrial function was the first biological process to be associated with PD ^8,9^.

An interest in mitochondrial function and PD began with the observation that exposure to the drug 1-methyl-4-phenyl-1,2,3,4-tetrahydropyridine (MPTP) can cause rapid parkinsonism and neuronal loss in the SN in humans, and that this is mediated through inhibition of complex I of the mitochondrial electron transport chain ^7,10,11^. Subsequent work suggested that individuals with sporadic PD have reduced complex I activity not only in the SN, but in other brain regions and peripheral tissues ^12^. Genetic studies focusing on monogenic forms of PD provided further support for the involvement of mitochondrial dysfunction in the disease. Pathogenic mutations that lead to autosomal recessive forms of PD have been reported in *PINK1, PARK2, PARK7, CHCHD2* and *VPS13C* and the proteins they encode are all now known to be involved in the mitochondrial quality control system and in particular mitophagy ^13–16^.

Therefore in this paper, we aim to comprehensively assess the role of mitochondrial function in sporadic PD by leveraging improvements in the scale and analysis of PD genome wide association study (GWAS) data with recent advances in our understanding of the genetics of mitochondrial disease. The availability of large scale genome wide association data in PD cases and the rapid identification of genetic lesions that underlie mitochondrial disease provide an opportunity to systematically assess the role of genetic variability in mitochondrial linked genes in the context of risk for PD^17^. In this study we combine these new resources with current statistical tools, such as polygenic risk scoring and Mendelian randomisation, to explore the role of mitochondrial function in both PD risk and age at onset of disease to obtain novel insights.

## RESULTS

### A component of the “missing heritability” of PD lies within mitochondria function genes

The general workflow for the genetic analysis used in this study is shown in **Figure 1.** First, to study the importance of mitochondrial function in sporadic PD, we investigated the heritability of PD specifically within genomic regions that contained genes annotated as important in mitochondrial function. The construction of this annotation was driven by the principle that genomic regions, which are known to be the sites of mutations in individuals with rare mitochondrial diseases or are candidate regions for such mutations provide the best evidence for involvement in mitochondrial function.

**Figure 1.**
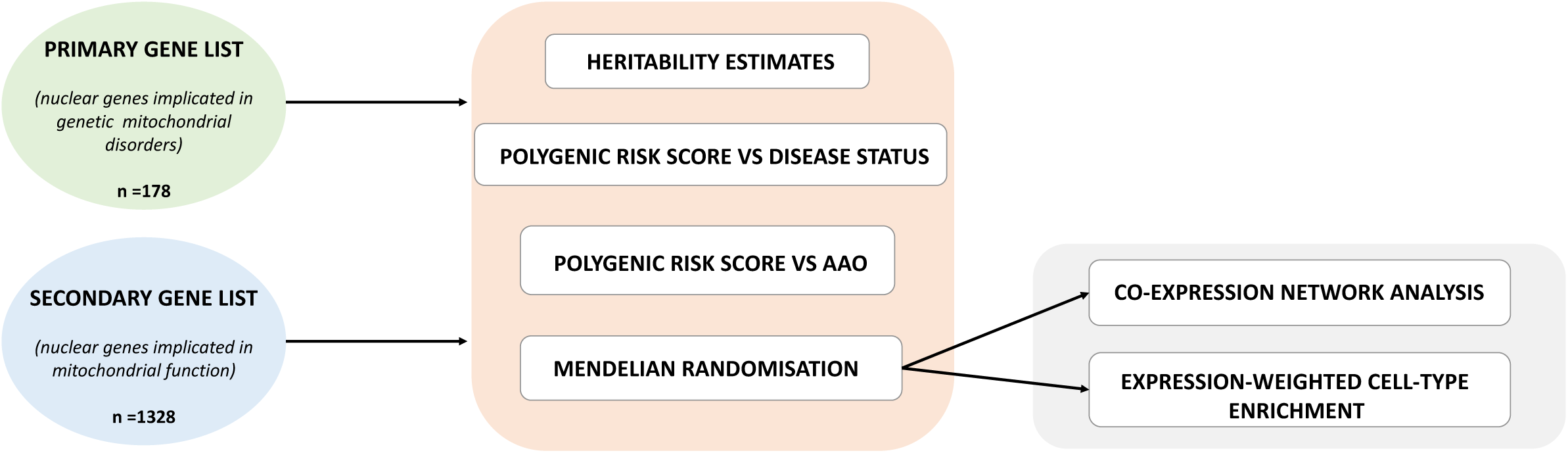
Workflow of mitochondrial-function specific PD analysis

Using GCTA, heritability estimates were first calculated for the four largest IPDGC GWAS datasets and including all variants (UK_GWAS, SPAIN3, NIA, DUTCH). Due to the low number of included cases, the heritability estimates in the other IPDGC datasets were deemed less reliable. Consistent with previous heritability estimates from both Keller and colleagues (2012; 24%) and Chang and colleagues (2017; 21%), our random effects meta-analysis for the four datasets identified 23% (95% CI 12-34, p= 2.72E-05) phenotypic variance associated with all PD samples (**Table 1 & 2**). There was a high degree of consistency across the cohorts.

**Table 1.** Cohort level heritability analysis for the primary and secondary mitochondrial gene lists

|  | Primary |  | Secondary |  |
| --- | --- | --- | --- | --- |
|  | Heritability estimate | SE of heritability estimate | Heritability estimate | SE of heritability estimate |
| UK_GWAS | 0.00321 | 0.00277 | 0.00563 | 0.00688 |
| SPAIN3 | 0.00027 | 0.00314 | 0.00629 | 0.00932 |
| NIA | 0.00945 | 0.00540 | 0.03616 | 0.01365 |
| DUTCH | 0.00000 | 0.00530 | 0.03562 | 0.01681 |

**Table 2.**
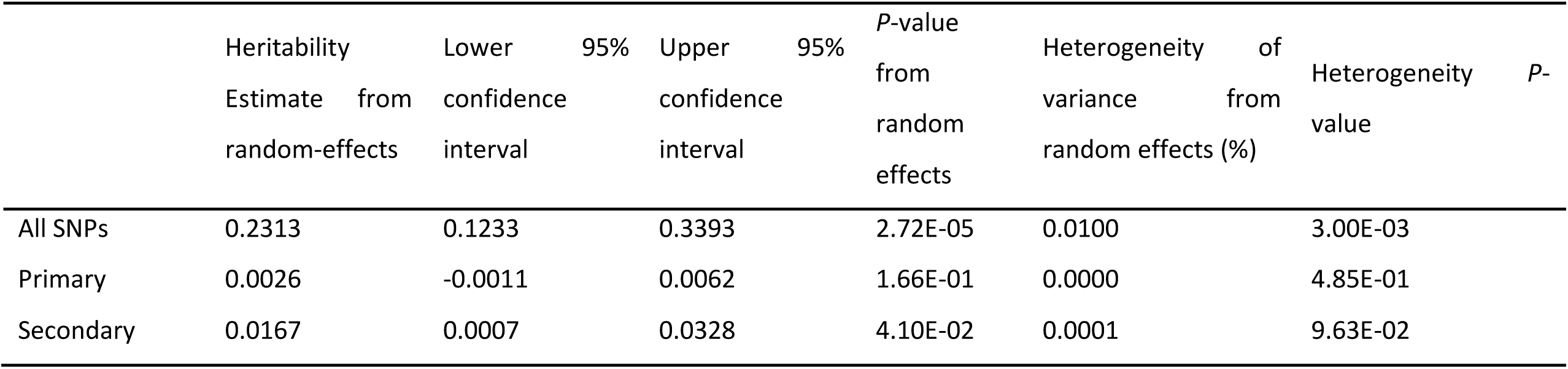
Summary of random-effects meta-analysis for the primary and secondary mitochondrial gene lists

After establishing the consistency of our heritability estimates we next calculated heritability using only variants located within genic regions specified as being of primary (n=176) or secondary (n=1463) importance in mitochondrial function. Genes within the primary or secondary lists, which had already been identified in the most recent PD meta-analysis were excluded ^6^. The heritability estimate using a random-effects meta-analysis for the primary gene list was estimated to be a modest 0.26% (95% CI −0.11-0.66, p=0.166). However, the heritability estimate using a random-effects meta-analysis for the secondary list, namely genes implicated in mitochondrial function or morphology as well as disease, was estimated to be 1.67% (95% CI −0.07-0.32, p=0.041).

### Mitochondria function specific polygenic risk score is significantly associated with disease status

Next, we calculated PRS to capture the addictive effect of all common variants within genes implicated in mitochondria function on PD risk. PRS is a particularly powerful approach in this context because it is able to efficiently incorporate information from all hits including sub-significant hits, which may nonetheless be etiologically relevant.

Using this approach the primary and secondary mitochondrial genomic annotations were found to be significantly associated with PD disease status. Remarkably, the primary gene list consisting of only 176 genes implicated in Mendelian mitochondrial disorders, was associated with PD with an odds ratio of 1.12 per standard deviation increase in the PRS from the population mean (random-effects p-value = 6.00E-04, beta = 0.11, SE = 0.03). The secondary gene list, which also included genes implicated in mitochondria function or morphology, was associated with PD with a higher odds ratio of 1.28 per standard deviation increase in the PRS from the population mean (random-effects p-value =1.9E-22, beta = 0.25, SE = 0.03) (**Figure 2**). Together, these analyses not only provide further support for importance of mitochondrial processes in PD, but potentially provide a tool for identifying PD patients most likely to benefit from treatments specifically targeting mitochondrial function.

**Figure 2.**
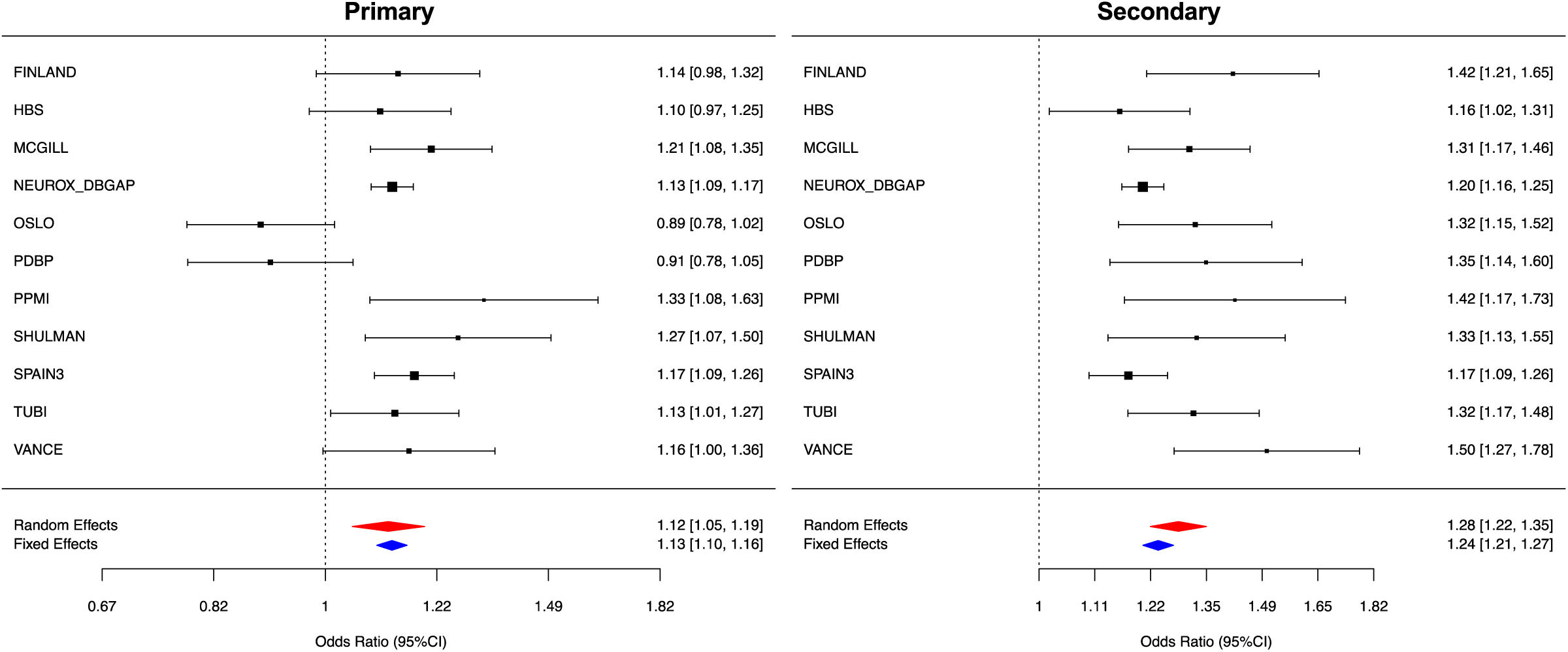
Forest plots of PRS for Parkinson’s Disease across cohorts. Random effect meta analysis results are shown as red diamonds and fixed effects are shown as blue, with the centerline of each diamond representing the summary PRS for that dataset.

### Mitochondria function-specific polygenic risk score is significantly associated with later age at onset

Although multiple lines of evidence point to the importance of mitochondrial dysfunction as a primary cause of PD, impaired mitochondrial dynamics appears to be common to a wide range of neurodegenerative diseases including Huntington’s disease^18,19^, amyotrophic lateral sclerosis^20,21^ and Alzheimer’s disease^22–25^. The latter suggests that even when impaired mitochondrial function is not the primary event in disease pathogenesis, it is a common outcome and could contribute to disease progression. We sought to test this hypothesis by investigating the importance of common variation within our mitochondrial gene lists in determining the age at onset of PD (AAO). Given the significant lag period between PD pathophysiology and symptoms, AAO was used as an indirect measure of disease progression. This analysis was performed using PRS since it has been consistently found to be the main genetic predictor of AAO ^6,26,27,28^ with higher genetic risk scores being significantly associated with an overall trend for earlier AAO of disease. While the primary mitochondrial gene list was not significantly associated with AAO of disease, the secondary gene list was correlated with AAO. Contrary to expectation, the cumulative burden of common variants within the 1463 genes comprising the PRS for PD risk, were positively correlated with AAO of PD. After meta-analysing all 11 cohorts, we found that each 1SD increase in PRS, led to a 0.62 year increase in the AAO of disease (summary effect = 0.618, 95%CI (0.325-0.911),|^2^=61.33%, p-value=3.56E-05, **Figure 3**). As the forest plots demonstrate, although there was a relatively high heterogeneity across studies, the directionality and magnitude of effect on AAO were in concordance with the meta-analysis with the exception of the Oslo cohort. This finding could suggest that firstly, disease causation and progression are genetically separable processes in PD and that secondly the role of mitochondrial dysfunction in PD is likely to be highly complex with multiple distinct mitochondrial processes likely to be involved at different disease stages.

**Figure 3.**
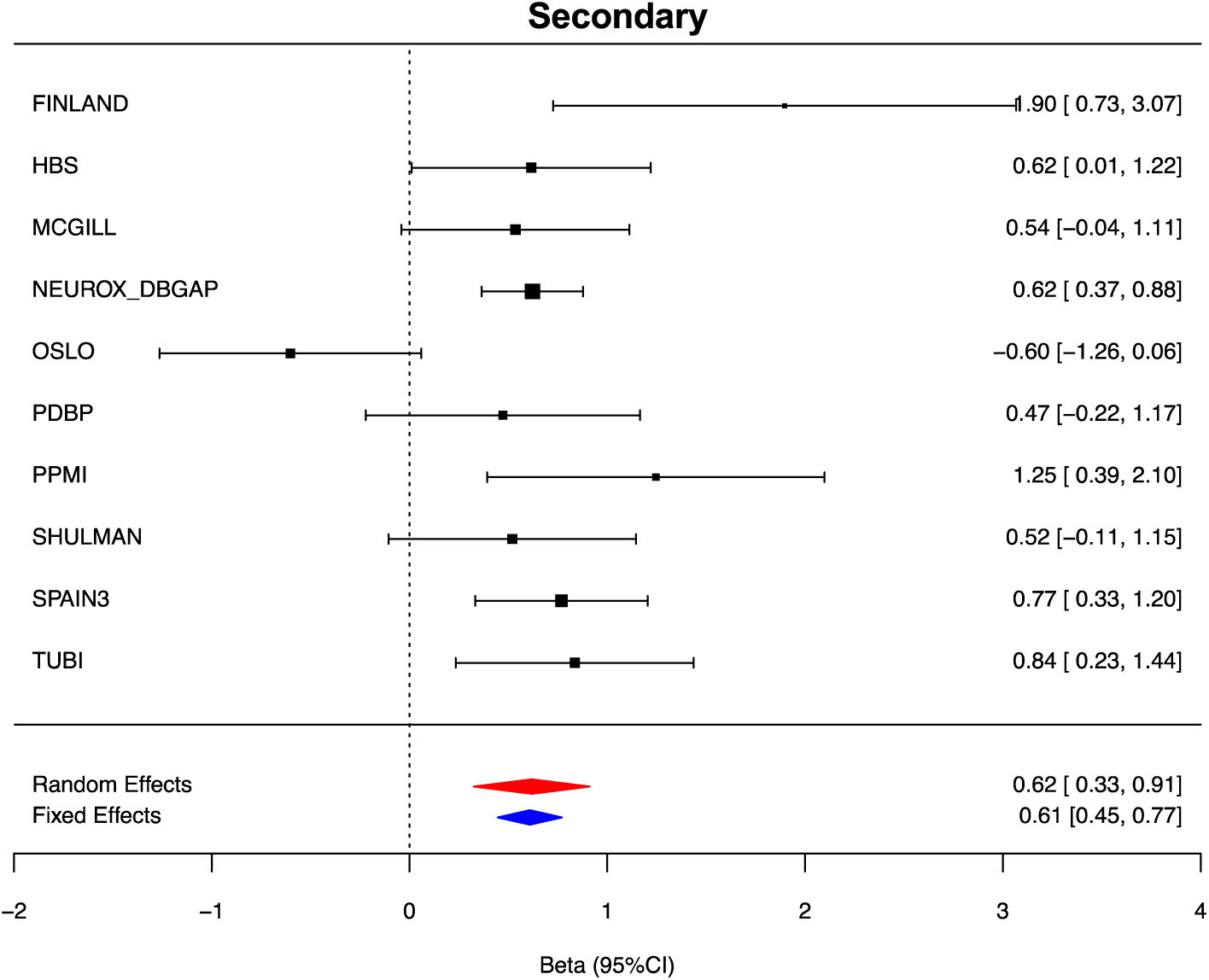
Forest plots of PRS for the age at onset of Parkinson’s Disease across cohorts. Random effect meta analysis results are shown as red diamonds and fixed effects are shown as blue, with the centerline of each diamond representing the summary PRS for that dataset.

### MR suggests potential causal association of fourteen novel mitochondria function genes with PD risk

Given the robust evidence for the involvement of mitochondrial function in sporadic PD, next we used two sample MR analysis to identify specific genes likely to be important in PD risk. Since we wanted to identify novel associations, we excluded genes already linked to PD through the most recent GWAS meta-analysis^6^. This resulted in the exclusion of 31 genes linked to mitochondrial function and in linkage disequilibrium with the top PD risk variants. Analysis of the remaining 1432 genes (generated by combining the primary and secondary gene lists) resulted in the identification of fourteen novel genes linked to mitochondrial function and causally associated with PD risk (**Table 3**). Of the fourteen genes, the expression of 5 genes (*CLN8, MPI, LGALS3, CAPRIN2* and *MUC1)* was positively associated with PD risk in blood. Similarly, in brain PD risk was associated with increased expression of *ATG14, E2F1*, and *EP300* in brain. However, negative associations in brain and blood expression were observed for *MRPS34* and *PTPN1 and LMBRD1* respectively. Finally, elevated methylation of *FASN* in the brain was found to be positively associated with PD risk but elevated methylation of *CRY2* was found to be inversely correlated.

**Table 3.**
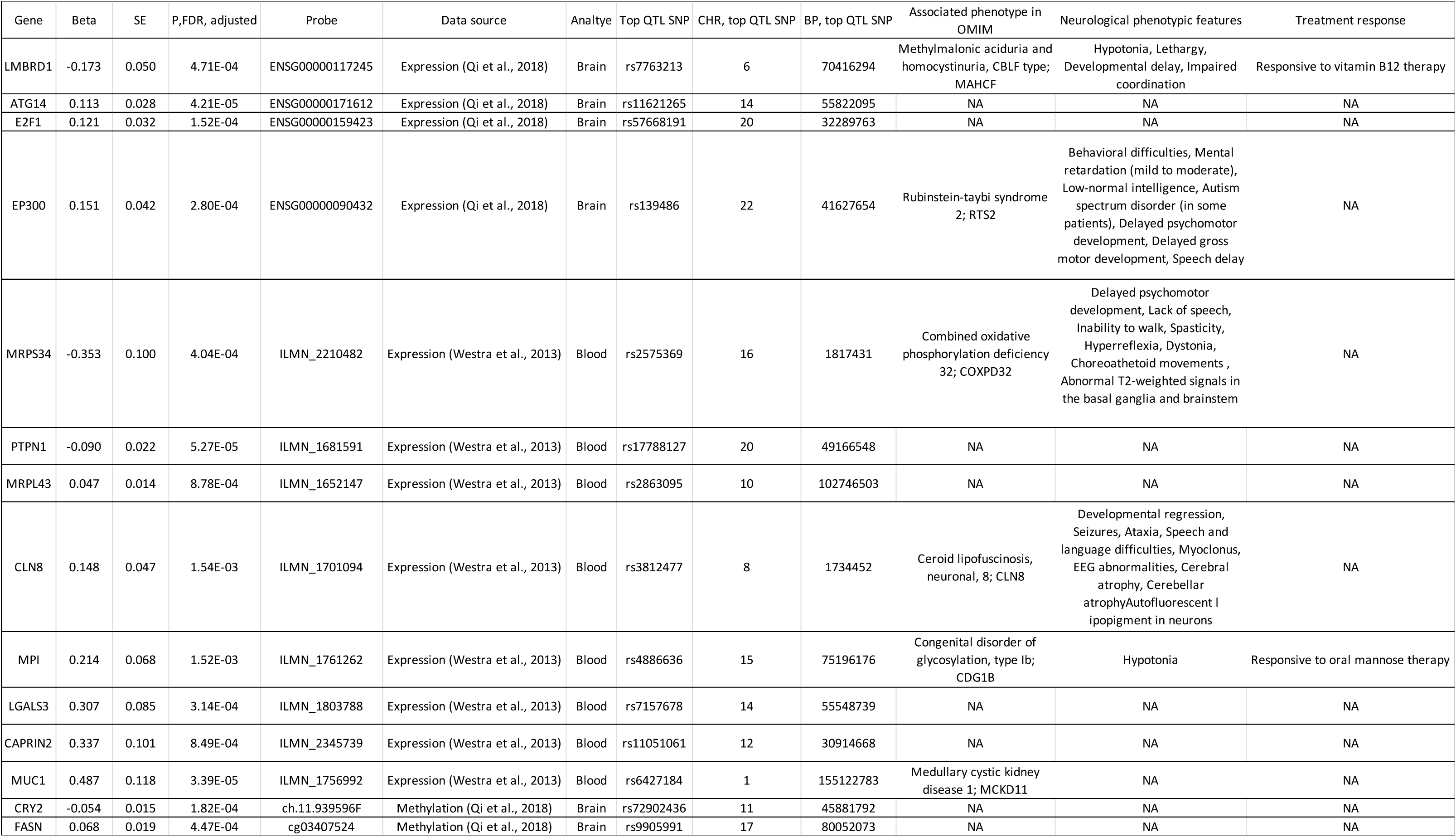
Significant functional associations of mitochondrial function genes via two-sample Mendelian randomization

Six of the fourteen novel PD risk genes we identified (*CLN8, EP300, LMBRD1, MPI, MRPS34* and *MUC1*) are already associated with a monogenic disorder (**Table 3**). We noted that neurological abnormalities were a feature of the condition in five of the six cases with Combined Oxidative Phosphorylation Deficiency 32 due to biallelic mutations in *MRPS34* being perhaps of particular interest. In common with PD, this condition is associated with abnormalities of movement, including dystonia and choreoathetoid movements. Mutations causing this condition result in decreased levels of *MRPS34* protein causing destabilisation of the small mitochondrial ribosome subunit and suggesting the involvement of mitochondrial processes distinct from mitophagy and mitochondrial homeostasis in PD. In this context, it is noteworthy that *MRPL43*, another nuclear gene encoding for a component of the large mitochondrial ribosome subunit is also highlighted by the MR analysis. Thus, this analysis not only enabled us to identify specific genes of interest, but also pointed to the role of multiple mitochondrial processes in PD distinct from mitophagy.

### Exploring the expression of the novel mitochondria risk genes provides additional support for their role in PD

We leveraged publicly available cell-specific and tissue-specific gene expression data to investigate the 14 mitochondria genes implicated in PD through MR. First, we used enrichment-weighted cell-type enrichment (EWCE) to determine whether the expression of mitochondrial PD-associated genes (as identified through MR and described above, n= 14) was enriched within a specific cell-type class or their subtypes. No significant enrichment of these genes was found in any of the tested neuronal and glial cell-type classes (**Supplementary Table 6, Supplementary Figure 1**). Next, we used co-expression network analysis to identify possible functional interactions between the 14 novel mitochondrial genes identified through MR and genes implicated in monogenic forms of PD. We found that five of the 14 genes assessed, *CLN8, FASN, MPI, MRPL43* and *MRPS3*, were highly co-expressed with at least one gene already implicated in monogenic PD in multiple brain regions (>3 brain regions, **Supplementary Table 7**). Interestingly, in the case of CLN8, *MRPL43* and *MRPS4*, our novel genes were co-expressed with a monogenic PD gene already implicated in mitochondrial function such as *PARK7*. Furthermore, with the exception of *CLN8*, (*FASN, MPI, MRPL43* and *MRPS3*), the novel mitochondrial gene was assigned to a co-expression module enriched for neuronal markers (**Table 4, Supplementary Table 8**).

## DISCUSSION

We first demonstrate that a proportion of the “missing heritability” of sporadic PD can be explained by additive common genetic variation within genes implicated in mitochondrial disease and function, even after exclusion of genes previously linked to PD through linkage disequilibrium with the top risk variants ^4–6,29–33^. In fact, using PRS, which efficiently incorporates information from sub-significant hits, we showed that cumulative small effect variants within only 196 genes linked to monogenic mitochondrial disease significantly associated with increased PD risk (with odds ratios of 1.12 per standard deviation increase in PRS from the population mean). These findings are important for two main reasons. Firstly, given that risk profiling performed in the recent PD meta-analysis did not identify a significant association with mitochondrial function ^4–6,29–33^ 17. Secondly, since the primary gene list consisted solely of the 196 genes mutated in monogenic mitochondrial disorders, this analysis highlights the increasingly close relationship between Mendelian and complex disease^7^.

In order to maximise the utility of this study, we used MR which identified 14 specific mitochondrial genes of interest with putative functional consequences in PD risk. We found that although a number of the genes we identified are clearly linked to known PD-related pathways, such as lysosomal dysfunction in the case of *CLN8* and *LMBRD1* or autophagy in the case of *ATG14*, others appeared to point towards new processes. In particular, this analysis highlighted the mitochondrial ribosome through the identification of the genes, *MRPL43* and *MRPS34*, encoding components of the large and small mitochondrial ribosome subunits respectively. Interestingly, biallelic mutations in *MRPS34* are known to cause a form of Leigh syndrome, characterised by neurodegeneration in infancy with dystonia and choreoathetoid movements due to basal ganglia dysfunction. Furthermore, we note that a recent study that utilized whole exome sequencing (WES) data from two PD cohorts to investigate rare variation in nuclear genes associated with distinct mitochondrial processes, not only provided support for the involvement of mitochondrial function in sporadic PD, but also identified the gene, *MRPL43,* which encodes a component of the large mitochondrial ribosomal subunit^34^. Interestingly, *MRPL43* and *MRPS34* were amongst five genes which were also highly co-expressed in human brain with genes already known to cause monogenic forms of PD. Whereas *MRPL43* and *MRPS34* were highly co-expressed with *PARK7* in modules enriched for neuronal markers, *FASN* and *MPI* were co-expressed with *ATP13A2,* and *CLN8* was located in modules containing *FBXO7* and enriched for oligodendrocyte markers. While this form of analysis does not provide information at the single cell level, it points to the possibility of pathway interactions between these gene sets. However, most importantly it implicates entirely distinct mitochondrial processes in PD risk.

Finally, and perhaps most remarkably using our mitochondrial gene lists we observe clear differences between disease causation and AAO in PD. Although PRS of the primary mitochondrial gene list was not significantly associated with AAO, the PRS of the secondary mitochondrial gene list was positively correlated (p value =3.6E-05), indicating association with later age at onset. However, given these findings it seems plausible that some mitochondrial processes may contribute to PD risk. Thus, this analysis is consistent with the findings of the most recent and largest AAO PD GWAS, which reported that not all the well-established risk loci are associated with AAO and suggested a different mechanisms for PD causation and AAO ^35^.

Although in this study we have comprehensively analyzed the largest PD datasets currently available with very specific and inclusive mitochondrial function gene lists, there are a number of limitations to our analyses. Firstly, there was a relative amount of heterogeneity in age at PD diagnosis within the AAO GWAS studies used. This was due to certain cohorts AAO being self-reported and other cohorts specifically recruiting younger onset cases. Nonetheless, the highly significant p-value we obtain for the association mitochondrial genes and AAO of PD (p-value=3.56E-05) and the recognized importance of mitochondrial function in aging would suggest that this finding is likely to be robust. Furthermore, it is important to recognize that our understanding of mitochondrial biology is far from complete and this is made evident by the fact that many individuals with probable genetic forms of mitochondrial disease remain undiagnosed. Finally, the statistical tools we have used in these analyses are currently limited. For example, MR ultimately relies on the availability of sufficient quantities of high quality eQTL data. However, as there is a future focus to; increase data-set sample size, report and characterize phenotypes such as AAO more accurately and continue to increase the number of identified mitochondrial disease and function genes, we will be able to further explore the role of specific mitochondrial processes in more detail and identify their distinct contribution to disease causation and progression.

In summary, in this study we provide robust evidence for the involvement of mitochondrial processes in sporadic PD, as opposed to its defined and well-established role in the monogenic forms of the disease. In relation to the 14 novel mitochondrial function genes that we have identified, our data suggests that it is not only mitochondrial quality control and homeostasis which contributes to PD risk but other key mitochondrial processes, such as the function of mitochondrial ribosomes, mirroring the biological complexity of mitochondrial disorders. Thus, this study opens the way for the identification of novel drug targets in PD causation and progression.

## METHODS

### Samples and quality control of IPDGC datasets

All genotyping data was obtained from IPDGC datasets, consisting of 41,321 individuals (18,869 cases and 22,452 controls) of European ancestry. Detailed demographic and clinical characteristics are given in **Supplementary Table 1** and are explained in further detail in along with detailed quality control (QC) methods^6,35^. For sample QC, in short, individuals with low call rate (<95%), discordance between genetic and reported sex, heterozygosity outliers (F statistic cut-off of > −0.15 and < 0.15) and ancestry outliers (+/- 6 standard deviations from means of eigenvectors 1 and 2 of the 1000 Genomes phase 3 CEU and TSI populations from principal components ^36^) were excluded. Further, for genotype QC, variants with a missingness rate of > 5%, minor allele frequency < 0.01, exhibiting Hardy-Weinberg Equilibrium (HWE) < 1E-5 and palindromic SNPs were excluded. Remaining samples were imputed using the Haplotype Reference Consortium (HRC) on the University of Michigan imputation server under default settings with Eagle v2.3 phasing based on reference panel HRC r1.1 2016^37,38^.

### Curation of genes implicated in mitochondrial disorders and associated with mitochondrial function

Gene lists were built to encompass different levels of evidence for involvement of the respective protein products in disease phenotypes that relate to mitochondrial function. The list of genes implicated in genetic mitochondrial disorders (“primary” gene list, n=196) has the most stringent criteria of evidence that the respective genes is related to mitochondrial dysfunction. It consists of 102 nuclear genes listed in MITOMAP (downloaded 2015) and 94 sourced from literature review as containing mutations that cause with mitochondrial disease.

The list of genes implicated in mitochondrial function (“secondary” gene list, n = 1487) was constructed using the OMIM API to identify all genes for which the word “mitochondria” (or derivatives) appeared in the free-text description, and by combining this information with MitoCarta v2.0 genes with no OMIM phenotype. This therefore gathered a list of plausible biological candidate genes, i.e. genes that are functionally implicated in mitochondrial function and morphology for which we may lack genetic evidence for mitochondrial disease association.

Next, to identify novel PD-associated genes, the 349 genes identified to be in LD with the PD risk variants of interest in the most recent PD meta-analysis (Nalls et al 2018) were removed from both lists (removed genes listed in **Supplementary Table 2**). The final “primary” and “secondary” gene lists are given in **Supplementary Table 3** and **Supplementary Table 4** and following the removal the PD-associated genes were n= 178 and n=1328 respectively.

### Cohort-level heritability estimates and meta analysis

Genome-wide complex trait analysis (GCTA) was used to calculate heritability estimates for the four largest IPDGC cohorts (UK_GWAS, SPAIN3, NIA, and DUTCH) using non-imputed genotyping data for all variants within both mitochondria gene lists using the same workflow as ^39^. GCTA is a statistical method that estimates phenotypic variance of complex traits explained by genome-wide SNPs, including those not associated with the trait in a GWAS. Genetic relationship matrices were calculated for each dataset to identify the genetic relationship between pairs of individuals. Genetic relationship matrices were then input into restricted maximum likelihood analyses to produce estimates of the proportion of phenotypic variance explained by the SNPs within each subset of data. Principal components (PCs) were generated for each data-set using PLINK (version 1.9). In order to adjust for factors accounting to possible population substructure, the top twenty generated eigenvectors from the PC analysis, age, sex and prevalence were used as basic covariates. Disease prevalence standardized for age and gender based on epidemiological reports was specified at 0.002^39–43^. Summary statistics from these estimates were produced for all four datasets and were included in the meta-analyses. Random-effects meta-analysis using the residual maximum likelihood method, was performed using R (version 3.5.1) package metafor to calculate p-values and generate forest plots^44^.

### Risk profiles versus disease status and age at onset

Previous risk profiling methods have calculated polygenic risk scores (PRS) using only variants that exhibit genome-wide significant associated with disease risk. However, in the most recent PD meta analysis, it is shown that using variants at thresholds below genome-wide significance improves genetic predictions of disease risk (^6,39^). Mirroring this workflow, but instead using only variants within gene regions outlined in both the primary and secondary gene lists, the R package PRsice2 was used to carry out PRS profiling in the standard weighted allele dose manner. In addition, PRsice2 performs permutation testing and p-value aware LD pruning to facilitate identifying the best p-value threshold for variant inclusion to construct the PRS. External summary statistics utilized in this phase of analysis included data from leave-one-out meta-analyses (LOOMAs) that exclude the study in which the PRS was being tested, avoiding overfitting/circularity to some degree. LD clumping was implemented under default settings (window size = 250kb, r^2^ > 0.1) and for each dataset 10,000 permutations of phenotype-swapping were used to generate empirical p-value estimates for each GWAS derived p-value threshold ranging from 5E-08 to 0.5, at a minimum increment of 5E-08. Each permutation test in each dataset provided a Nagelkerke’s pseudo r^2^ after adjustment for an estimated prevalence of 0.005 and study-specific PCs 1-5, age and sex as covariates. GWAS derived p-value threshold with the highest pseudo r^2^ was selected for further analysis. Summary statistics were meta-analyzed using random effects (REML) per study-specific dataset using PRSice2 ^45^. For the age at onset risk profiling, the same workflow was followed, however instead, age at onset was used as a continuous variable, as previously reported^35^

### Mendelian randomization to explore possible causal effect of mitochondria function genes

MR uses genetic variants to identify if an observed association between a risk factor and an outcome is consistent with causal effect ^46^. This method has been implemented in several recent genetic studies to identify association between expression quantitative trait loci (QTL), to more accurately nominate candidate genes within risk loci. Therefore, for this study, in the aim of identifying whether changes in expression of mitochondria function genes are potentially causally related to PD risk, two-sample MR was implemented. Both mitochondria gene lists were combined and all genes already associated with PD (i.e. that have been identified to be in LD with PD risk loci in the last meta-analysis) were removed, leaving 1432 unique mitochondria function gene regions.

We utilized four large-scale methylation and expression datasets through the summary-data-based Mendelian randomization (SMR) (http://cnsgenomics.com/software/smr) resource. Summary statistics were compared to PD outcome summary statistics for the mitochondria variants of interest (extracted from ^4–6,29–33^) to identify possible association using the R package TwoSampleMR.

Tissues were selected based on their relevance to the pathobiology of PD, which ultimately consisted of tissues from 10 brain regions, whole blood, skeletal muscle, and nerve, (a full list of the tissues utilised can be found in **Supplementary Table 5**). For the methylation QTLs “middle age” data was used, which was the the oldest available time point. For each mitochondria function variant of interest (considered here the instrumental variable), wald ratios were generated for each variable that tagged a cis-QTL (probes within each gene and meeting a QTL p-value of at least 5E-08 in the original QTL study) and for a methylation or expression probe with a nearby gene. Using the default *SMR* protocols, linkage pruning, and clumping were implemented. Finally, for each dataset p-values were adjusted by false discovery rate to account for multiple testing.

### Co-expression network analysis

Co-expression network analysis was used to determine whether mitochondrial genes associated with PD using the SMR analysis are co-expressed with genes associated with monogenic forms of PD in human brain. This analysis was performed by using GTEx V6 gene expression data (The GTEx Consortium et al. 2015; Carithers et al. 2015) to generate co-expression networks for each of the 13 brain tissues included within the GTEx study. The raw FPKM (Fragments Per Kilobase of transcript per Million mapped reads) values were corrected for known batch effects, age at death, sex and post-mortem interval, as well as unknown effects. The unknown effects were detected with the Surrogate Variable Analysis (SVA) R Package (Leek and Storey 2007) and correction was performed using ComBat (Johnson, Li, and Rabinovic 2007). The resulting residuals were used to create a signed network using the blockwiseConsensusModules R function from the WGCNA R package (Langfelder and Horvath 2008) for each of the 13 tissues. Next, the modules obtained in each of the 13 networks were assigned to cell types using the userListEnrichment R function implemented in the WGCNA R package, which measures enrichment between module-assigned genes and defined brain-related lists using a hypergeometric test. The same approach was used to annotate modules with Gene Ontology, REACTOME (Fabregat et al. 2018) and KEGG (Kanehisa et al. 2016) terms.

### Expression-weighted cell-type enrichment (EWCE): evaluating enrichment of mitochondrial genes associated with PD risk

Expression-weighted cell-type enrichment (EWCE) (https://github.com/NathanSkene/EWCE) (EWCE) was used to determine whether mitochondrial genes associated with PD using the MR analysis have higher expression within a particular cell type than expected by chance. The input for the analysis was 1) neuronal and glial clusters of the central nervous system (CNS) identified in the Linnarsson single-cell RNA sequencing dataset (amounting to a subset of 114 of the original 265 clusters identified) (http://mousebrain.org/) and 2) our list of mitochondrial genes highlighted through the MR analysis (see **Supplementary Table 6** for the full list of Linnarsson CNS neuronal clusters used).

For each gene in the Linnarsson dataset, cell-type specificity was estimated (the proportion of a gene’s total expression in one cell type compared to all cell types) using the ‘generate.celltype.data’ function of the EWCE package. EWCE with the target list was run with 100,000 bootstrap replicates, which were sampled from a background list of genes that excluded all genes without a 1:1 mouse: human ortholog. In addition, transcript length and GC-content biases were controlled for by selecting bootstrap lists with comparable properties to the target list. The analysis was performed using major cell-type classes (e.g. “telencephalon inhibitory interneurons”, “telencephalon projecting excitatory neurons”, etc.) and subtypes of these classes (e.g. TEINH6 [“Interneuron-selective interneurons, cortex/hippocampus”], TEINH7 [“Interneuron-selective interneurons, hippocampus”], etc.). Data are displayed as standard deviations from the mean, and any values < 0, which reflect a depletion of expression, are displayed as 0. P-values were corrected for multiple testing using the Benjamini-Hochberg method over all cell types and gene lists displayed.

### OMIM data

Phenotype relationships and clinical synopses of all OMIM genes were downloaded from http://api.omim.org on the 29th of May 2018. OMIM genes were filtered to exclude provisional, non-disease and susceptibility phenotypes retaining 2,898 unique genes that were confidently associated to 4,034 Mendelian diseases. The phenotypic information relating to all genes associated with mitochondrial disorders was collated.

## Supporting information

## ACKNOWLEDGEMENTS

We would like to thank all of the subjects who donated their time and biological samples to be a part of this study. We also would like to thank all members of the International Parkinson Disease Genomics Consortium (IPDGC). See for a complete overview of members, acknowledgements and funding http://pdgenetics.org/partners. This work was supported in part by the Intramural Research Program of the National Institute on Aging, National Institutes of Health, Department of Health and Human Services; project ZO1 AG000949 and The Lily Foundation UK. MR was supported by the UK Medical Research Council (MRC) through the award of Tenure-track Clinician Scientist Fellowship (MR/N008324/1). John Hardy’s contribution was in part supported by MR/N026004/1. A full acknowledgement for the data-sets used for this work is available in the Supplementary data of the *Nalls et al* 2018^6^.

## AUTHOR CONTRIBUTIONS

### Data generation and analysis

KJB, IAB, SB-C, CD, JAB, RHR, DZ, MAS, CB, ZG-O, JRB, MAN, AS, MR, SK

### Design and funding

KJB, IAB, SB-C, JQ, VJB, CD, JAB, RHR, MAS, JRB, MAN, AS, MR, SK

### Critical review and writing the manuscript

KJB, IAB, SB-C, JQ, VJB, CD, JAB, RHR, MAS, CB, JRB, ZG-O, MAN, AS, MR, SK

## SUPPLEMENTARY

**Table S1.** Demographic and clinical characteristics for all IPDGC genotyping data

**Table S2.** Mitochondrial genes identified to be in LD with the PD risk variants of interest in the most recent PD meta-analysis that were removed from our list to identify novel association

**Table S3.** Primary mitochondria gene list (hg/19)

**Table S4.** Secondary mitochondria gene list (hg/19)

**Table S5.** Tissues and data-sets included in Mendelian Randomisation analysis

**Table S6.** List of the CNS cell clusters used for enrichment-weighted cell-type enrichment (EWCE) and results of EWCE analysis for all genes identified by two-sample Mendelian randomization

**Table S7** Coexpression of genes identified using two-sample Mendelian randomization with genes known to cause familial forms of PD

**Table S8.** Results of co-expression network analysis for all genes identified by two-sample Mendelian randomization

**Figure S1.** Results of enrichment-weighted cell-type enrichment analysis for all genes identified by via two-sample Mendelian randomization

