## Supplementary figures and images for "Genetic variation within genes associated with mitochondrial function is significantly associated with later age at onset of Parkinson disease and contributes to disease risk"

### Supplementary file 2

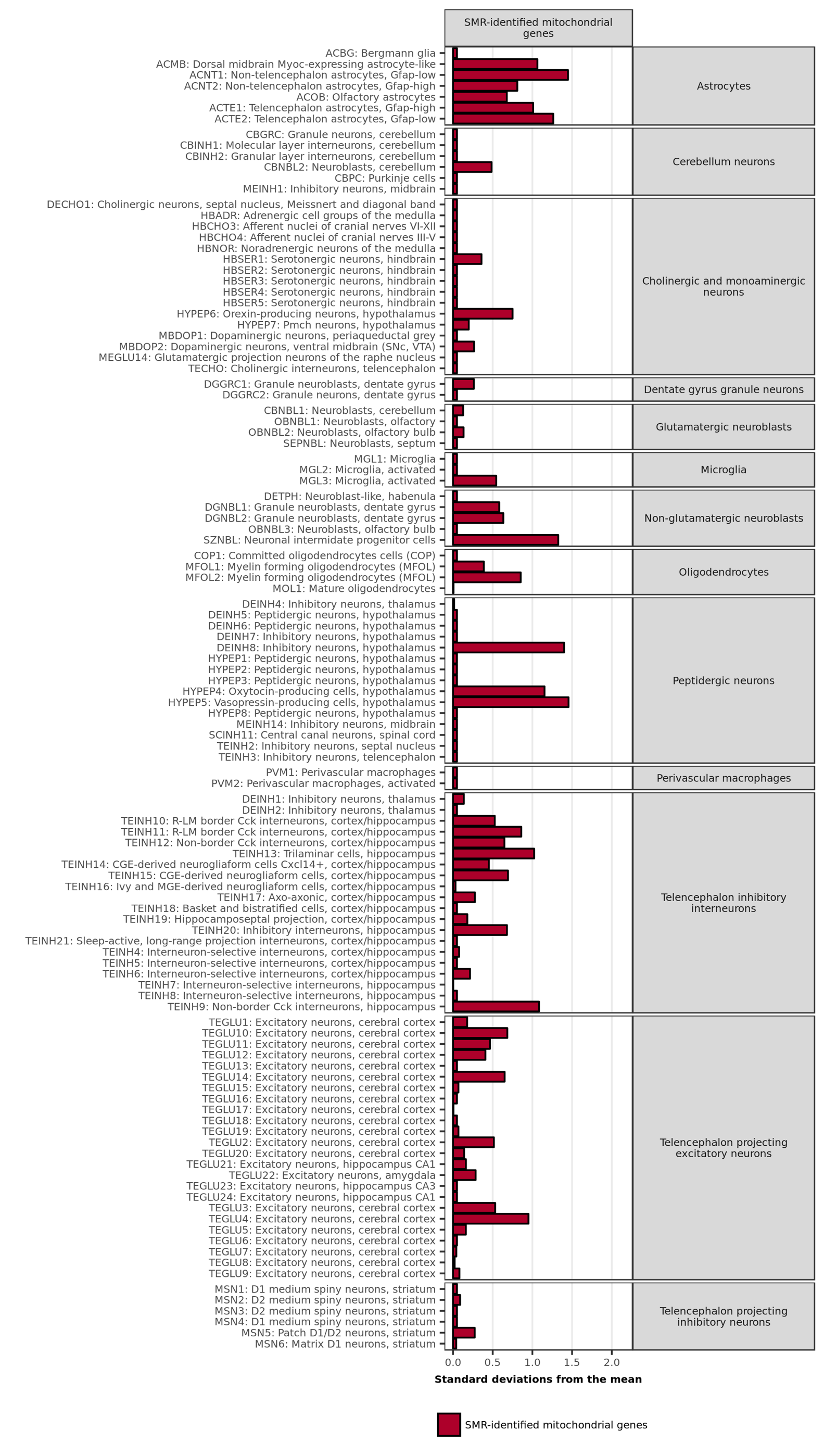
